# Characterization of cotton virus A, a novel and distinct member of the genus *Caulimovirus* with endogenous viral elements in *Gossypium* spp

**DOI:** 10.1101/2023.06.14.544975

**Authors:** Michael West-Ortiz, Douglas Stuehler, Emma Pollock, Jennifer R. Wilson, Stephanie Preising, Adriana Larrea-Sarmiento, Olufemi Alabi, Marc Fuchs, Michelle Heck, Alejandro Olmedo-Velarde

## Abstract

The complete genome sequence of cotton virus A, a new virus infecting cotton (*Gossypium* spp.), was determined using high-throughput sequencing, PCR and rolling circle amplification. The 7,482-nt genome is a circular dsDNA molecule that codes for six open reading frames similar to other members of the *Caulimovirus* genus (family *Caulimoviridae*) but presenting key genome organization differences. P3, P5 and P6 are presumed movement, coat, and reverse transcriptase proteins, respectively. P1, P2 and P4 showed no homology to virus proteins. Endogenous viral elements were found in *G. hirsutum* and *G. tomentosum*, but not in *G. barbadense, G. arboreum,* or *G. herbaceum.*

---

Cotton (*Gossypium spp.*) is one of the world’s most important natural fibers. The cotton industry’s global, yearly economic impact has been estimated to be more than $600 billion dollars [1]. Cotton leafroll dwarf virus (CLRDV) is an emerging viral pathogen of cotton that is transmitted by the cotton aphid, *Aphis gossypii*, and the causative agent of cotton blue disease in South America [2], and cotton leafroll dwarf disease in the USA [3-6]. Symptoms, including stunting, leaf rolling, interveinal chlorosis, bronzing and reduced boll sets, vary depending on biotic and abiotic factors such as cultivar, environment, and stage of growth, and can be aggravated by other underlying stress [3]. The diversity of symptoms and incidence of asymptomatic, CLRDV-infected plants in the USA highlights a need to clarify if there are additional viral agents in the field co-infecting cotton with CLRDV that alter disease progress. Therefore, we characterized virus populations associated with CLRDV-infected plants in Mississippi.

In 2019, one *G. barbadense* and nine *G. hirsutum* plants displaying leaf rolling and distortion were uprooted and collected from Stoneville, MS and hand-carried to Cornell University in Ithaca, NY under USDA-APHIS-PPQ permit # P526P-22-06971. Once at Cornell, the plants were potted in 2-gallon pots containing 50% Cornell soil mix and 50% coarse perlite, and maintained in insect-proof cages in the greenhouse with bi-weekly fertilization, pest management and pruning as needed. In 2022, leaf midrib and petiole samples were collected from each plant. Additionally in 2022, six *Gossypium* spp. samples comprising leaf midribs and petioles were also collected from plants growing at Stoneville, MS, and displaying leaf distortion, bronzing, and interveinal chlorosis. Total RNA was extracted from all samples using the Spectrum™ Total RNA Kit (Sigma-Aldrich, USA), as per the manufacturer’s instructions. Three composite samples were prepared by pooling 2, 3 and 5 total RNA samples, respectively, using the 2019 samples, whereas the 2022 samples were pooled into a fourth composite sample. After depleting ribosomal RNA from the composite samples using the RiboMinus Plant kit for RNA-seq (ThermoFisher Scientific), cDNA libraries were constructed using the TruSeq RNA library prep kit (Illumina). The libraries were sequenced on an Illumina® NovaSeq 6000 system to obtain paired-end reads (2 × 100 bp) at the Genomics High-Throughput Sequencing (HTS) Facility at the University of California, Irvine.

The data obtained from ∼ 66-78 million raw reads per composite sample were processed following previously described workflow [7]. The identity of contigs larger than 500 bp was annotated by BLASTx searches of the NCBI virus sequence database as of September 2022. The annotated virus contigs revealed sequence similarity to CLRDV and caulimoviruses. The four caulimovirus contigs (2,043 to 7,217 nt), one from each of the four composite samples, shared ∼98% nucleotide identity with each other, had significant matches with dahlia mosaic virus (DMV) and other caulimoviruses, but were sufficiently divergent suggesting they represented a sequence of a potential new virus within the family *Caulimoviridae*.

Since some caulimovirus genomes have been found integrated into their host genome and regarded as endogenous viral elements (EVEs) [8], DNase-RT-PCR assays were performed with contig-specific primers (Table S1) on DNase-treated RNA using all the 2019 and 2022 samples, as previously detailed [9] to detect viral RNA and exclude amplification of DNA endogenous viral elements. This assay included RNA extracted from an uninfected *G. hirsutum* seedling as a negative control. Caulimovirus-like transcripts were detected in all the ten 2019, and four 2022 samples, except for the negative control (Figure 1A). Sanger sequencing of purified PCR amplicons revealed they had 100% nucleotide identity with the Illumina sequence contigs. Furthermore, rolling circle amplification (RCA) was used to amplify the episomal circular DNA virus genome and distinguish it from possible EVEs in *Gossypium*. Total nucleic acids were extracted using the OPS Synergy 2.0 Plant DNA Extraction Kit (OPS Diagnostics, Lebanon, NJ, USA), and subjected to RCA using Equiphi29 DNA polymerase (Thermoscientific, Whatman, PA), as per the manufacturer’s instructions. RCA reactions were digested with *KpnI*. A ∼2.1 kbp DNA fragment was observed by agarose gel electrophoresis for all samples (data not shown). The amplicons were gel-purified and directly sequenced at PlasmidSaurus (Eugene, OR) using Oxford Nanopore sequencing. The consensus sequence (30x coverage) showed 99% nucleotide identity to the caulimovirus Illumina contig sequence, confirming the episomal presence of a putative new caulimovirus in the cotton samples.

**Figure 1.**
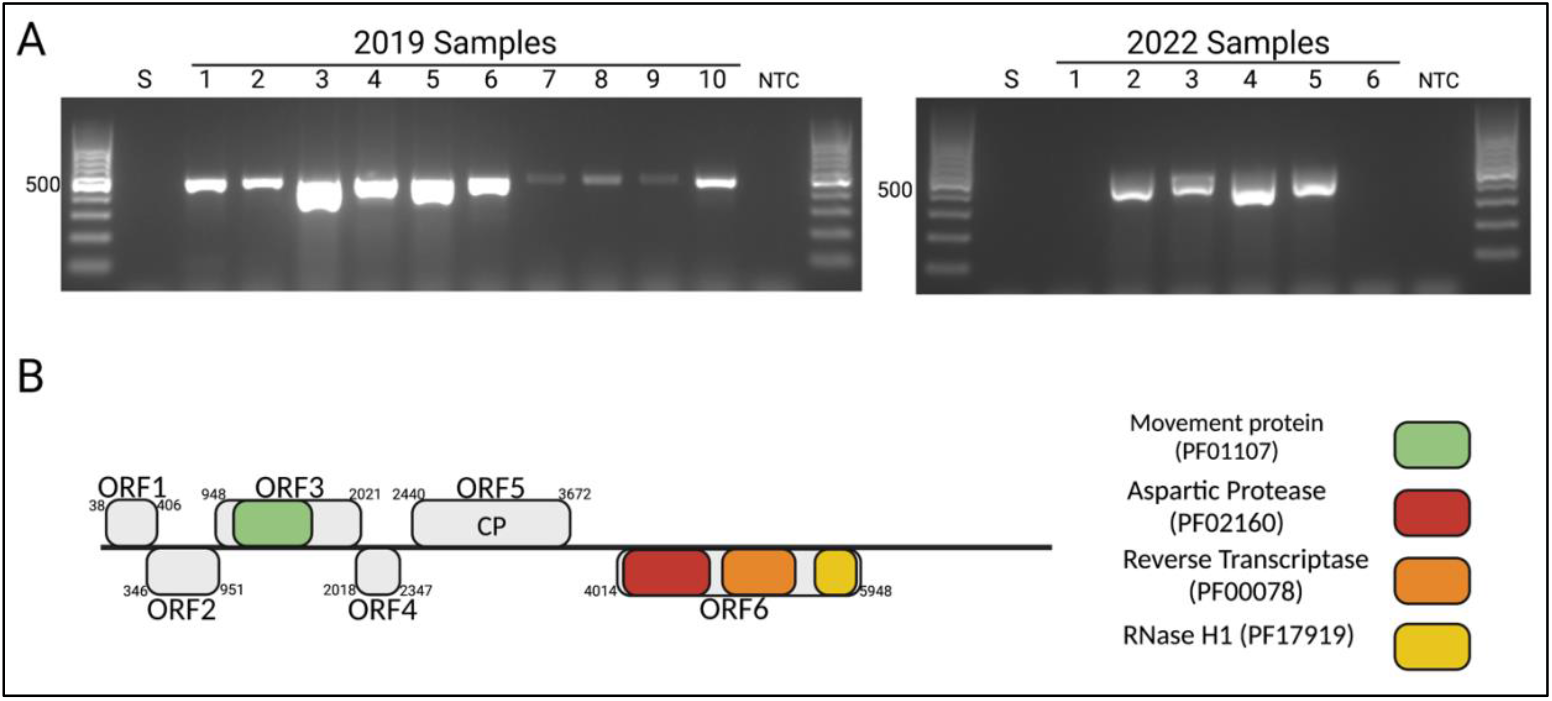
Detection of virus transcripts using DNase-RT-PCR assays (A) and a cartoon of the genome organization of cotton virus A (CotV-A) (B). A) RNA extracts of 2019 (lanes 1-10) and 2022 (lanes 1-6) samples were DNase-digested and subjected to RT-PCR using movement protein (MP) gene-specific primers to amplify a 474 bp product (Table S1). S = *Gossypium hirsutum* seedling. NTC = non-template control. B) A cartoon of the genome organization and predicted open reading frames (ORF) coding for six putative proteins that have conserved virus protein domains: MP (PF01107), aspartic protease (PF02160), reverse transcriptase (PF00078) and RNase H1 (PF17919). The genome is drawn to scale.

To obtain the complete genome sequence of the novel virus and validate the HTS results, one 2022 sample was selected for further analysis and five contig-specific overlapping primer sets (Table S1) were designed to fill gaps in the circular genome of the virus. PCR assays were performed using ten-fold diluted RCA products as the template. The amplicons were cloned, and three clones per amplicon were sequenced using Sanger and Nanopore sequencing. The complete genome of the novel virus comprises a 7,482-nt dsDNA circular molecule (GenBank accession). We consider the sequence to represent a new caulimovirus for which the common name cotton virus A (CotV-A) and species name Caulimovirus gossypii are proposed.

The genome of CotV-A starts with the conserved tRNAMet sequence 5′-TGGTATCAGAGCC-3′ and shares 66.7% nucleotide sequence identity with grapevine pararetrovirus (OP886324) with 22% query coverage. Six putative ORFs (>300 bp) were identified in the complete CotV-A genome sequence (Figure 1B) using ORF Finder (https://www.ncbi.nlm.nih.gov/orffinder/). ORF1 is at positions 38-406 nt; ORF2, 346-951 nt; ORF3, 948-2021 nt; ORF4, 2018-2347 nt; ORF5, 2440-3672 nt; and ORF6,4014-5948 nt. The predicted amino acid sequences of the putative proteins (P1 to P6) encoded by ORF 1-6 were searched against the Pfam and GenBank databases using the InterPro and BLASTP servers, respectively, for functional annotation of the conserved domains/motifs. P1 (122 aa, 14.4 kDa), P2 (201 aa, 24.4 kDa) and P4 (109 aa, 12.8 kDa) did not display homology to any known caulimovirus or known virus proteins. In contrast, P3 (357 aa, 41 kDa) represents a putative movement protein (MP) (PF01107), P5 is a putative viral coat protein (CP) (410 aa, 49.5 kDa) and P6 (644 aa, 75.9 kDa) is a typical polyprotein, akin other caulimoviruses, containing aspartic protease (PF02160, aa 22-229), reverse transcriptase (RT) (PF00078, aa 285-470) and RNase H1 (PF17919, aa 534-629) motifs. CotV-A has three intergenic regions (IGRs) (Figure 1B); the shorter IGR is 92 bp in length separating ORFs 4 and 5, the intermediate IGR is 341 bp and is between ORFs 5 and 6, whereas the long IGR (1571 bp) lies between ORFs 1 and 6. In the intermediate and long IGR, tentative TATA-like boxes were found in addition to a polyadenylation signal in the large IGR, which is a feature typical of caulimoviruses [10]. The nonanucleotide ‘TATAAAAAA’ (nt 3859-3867) was found in the intermediate IGR, and ‘TATAAAT’ (nts 6683-6689, 7264-7270, 7395-7401), ‘TATAAAAT’ (nt 6997-7004) and the polyadenylation signal ‘AATAA’ (nt 7472-7476) were found in the long IGR.

Pairwise sequence comparisons showed that the overall nucleotide sequence identity between CotV-A and other caulimoviruses ranged from 62.92% to 66.74%. The amino acid sequence identities of the CotV-A putative P3, P5 and P6 with homologs of other caulimoviruses ranged from 23.15% to 54.40% (Table S2). To gain insight into the taxonomic position of CotV-A and its relatedness to members of the family *Caulimoviridae*, phylogenetic analysis using the maximum-likelihood algorithm based on the RT-RNaseH protein region sequence was performed using the best model of protein evolution (rtREV+G+I) in MEGA 11 v11.0.13 [11] after 1,000 bootstrap replicates. The results showed the clustering of CotV-A into the *Caulimovirus* genus clade in a subclade also grouping grapevine pararetrovirus and plant-associated caulimovirus 1 (Figure 2). For members within the genus *Caulimovirus*, the species demarcation criterion requires divergence of more than 20% in the RT-RNase nucleotide region. Using pairwise comparisons, the sequence identities of this nucleotide region of CotV-A and caulimovirus homologs were all below 66.6%. This result provides evidence that CotV-A is a putative new species in the genus *Caulimovirus*.

**Figure 2.**
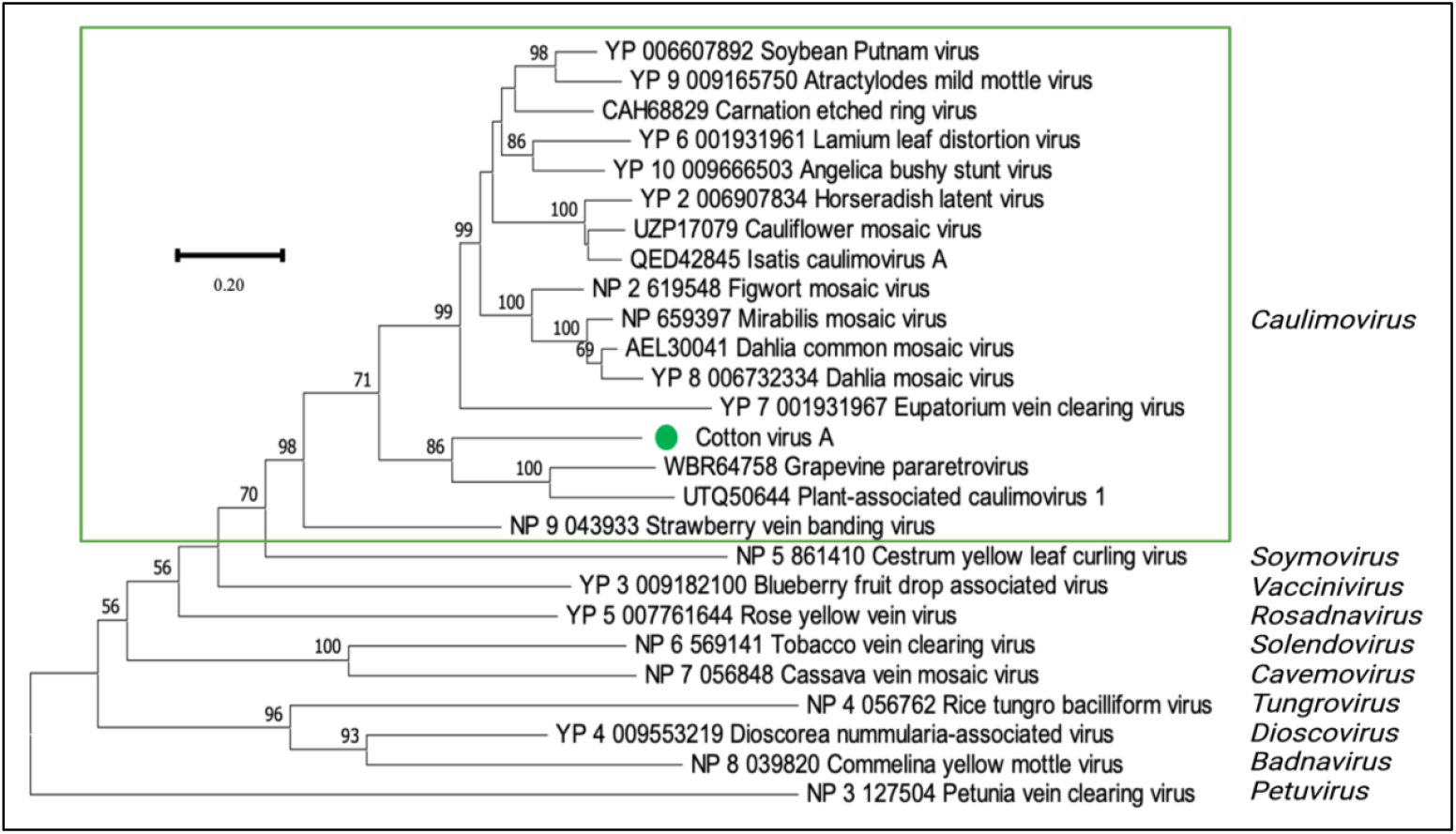
Phylogenetic relationships of the RT-RNase H protein sequence of cotton virus A (CotV-A) in relation to other members of the family *Caulimoviridae*. The maximum-likelihood method with the best model of protein evolution (rtREV+G+I) was used with 1,000 bootstrap replicates as percentage values for branch support. Predicted amino acid sequences were used and the respective GenBank accession number is shown with each virus name. The alignment was generated using MUSCLE and implemented in MEGA v.11.0.13. The green circle indicates CotV-A.

To evaluate CotV-A integration into the cotton genome, BLASTn searches were performed using the CotV-A genome sequence as query against chromosome-level genome sequences of the four commercially grown *Gossypium* species, *G. hirsutum, G. barbadense, G. arboreum* and *G. herbaceum.* Also, chromosome-level genome sequences of *G. tomentosum*, a very close wild *Gossypium* species was analyzed. All the *Gossypium* spp. genome sequences used in this study were from the CottonGen database (www.cottongen.org/) [12]. Additionally, mapping of quality trimmed virus reads was performed against the *Gossypium* spp. genome sequences using the Geneious mapper [13]. A total of four structurally unique EVEs were found in *G. hirsutum* and *G. tomentosum*. These contained the same predicted conserved protein domains as CotV-A (Figure 3A). No EVE was found in the other *Gossypium* species. In *G. hirsutum*, chromosomes A04 (11 out of the 13 genomes analyzed) and D12 (1 out of the 13 genomes analyzed) contained EVEs as inverted tandem repeat sequences (EVE 1 and 2, Figure 3B) which presented ∼99% nucleotide identity, and 84.5-100% protein identities to CotV-A (Table S3). Also, a single integration event was found in chromosome D03 (13 out of the 13 genomes analyzed) which was not only truncated (EVE 3, Figure 3B) but also presented ∼80% nucleotide identity, and 54-94% protein identities to CotV-A. Interestingly, EVE 1, which is either inserted in the sense direction (chromosome A04), or in the anti-sense direction (chromosome D12), contains two additional ORFs which were not present in the episomal CotV-A genome (Figure 3B). BLASTx searches of the region containing both ORFs showed that the concatenated predicted protein sequence shares low (<27%) identity to the P6 of grapevine pararetrovirus and other caulimoviruses. P6 is a multifunctional protein involved in viroplasm formation, and other virus cycle processes as well as suppression of antiviral defenses [14] In EVE 2, in each tandem repeat on chromosomes A4 and D12, the CP and the polyprotein sequences were interrupted, and spread across two predicted ORFs (Figure 3A and 3B). *In G. tomentosum*, chromosomes A03 (2 out of 2 genomes analyzed) contained a single virus integration event (EVE 4) which lacks ORF 1 and 2, and similar to EVE 2, the CP and the polyprotein sequences are each coded across two ORFs (Figure 3A and 3B). EVE 4 presented ∼79% nucleotide identity, and 71-89% protein identities to CotV-A (Table S3). The genomic locations in the chromosomes where EVE 1-3 are inserted were analyzed in the *G. hirsutum* genomes described above using bedtools v2.27.1 [15] and chromoMap v4.1.1 [16]. All predicted proteins 500 kb up- and downstream of each EVE were aligned to the KOfam database via KofamKOALA [17] with an e-value cutoff of 1e-5. Of the A4 and D12 integration sites (Figure S2A), adjacent proteins were found to belong to metabolic pathways (map01100), aminoacyl-tRNA biosynthesis (map00970), and plant hormone signal transduction (map04075) pathways. Coding sequences (CDS) surrounding the truncated D3 integration site (Figure S2B) were found to encode proteins belonging to biosynthesis of secondary metabolites (map01110), glycolysis/gluconeogenesis (map00010), ABC transporters (map02010), zeatin biosynthesis (map00908), and phenylpropanoid biosynthesis (map00940) pathways. Averaging 19.4 kb upstream of the A4 and D12 sense direction integration sites (Figure S2A), all 12 genomes encode an amino-acid permease BAT1 of 549 amino acids, responsible for amino acid transport. Downstream of the CotV-A anti-sense direction integration, averaging 15.9 kb between 11 samples, lies a CDS encoding a putative triacylglycerol lipase. The only genome which did not have triacylglycerol lipase adjacent to its anti-sense CotV-A integration was TX1000, where the protein was found to be encoded ∼30 Mbp upstream in chromosome A4. At all D3 integration sites the 5’ partial CotV-A virus genome is ∼3 kb away from the Piezo type mechanosensitive ion channel component 2 CDS, and the 3’ end is ∼12 kb away from the CDS of O-fucosyltransferase 39.

**Figure 3.**
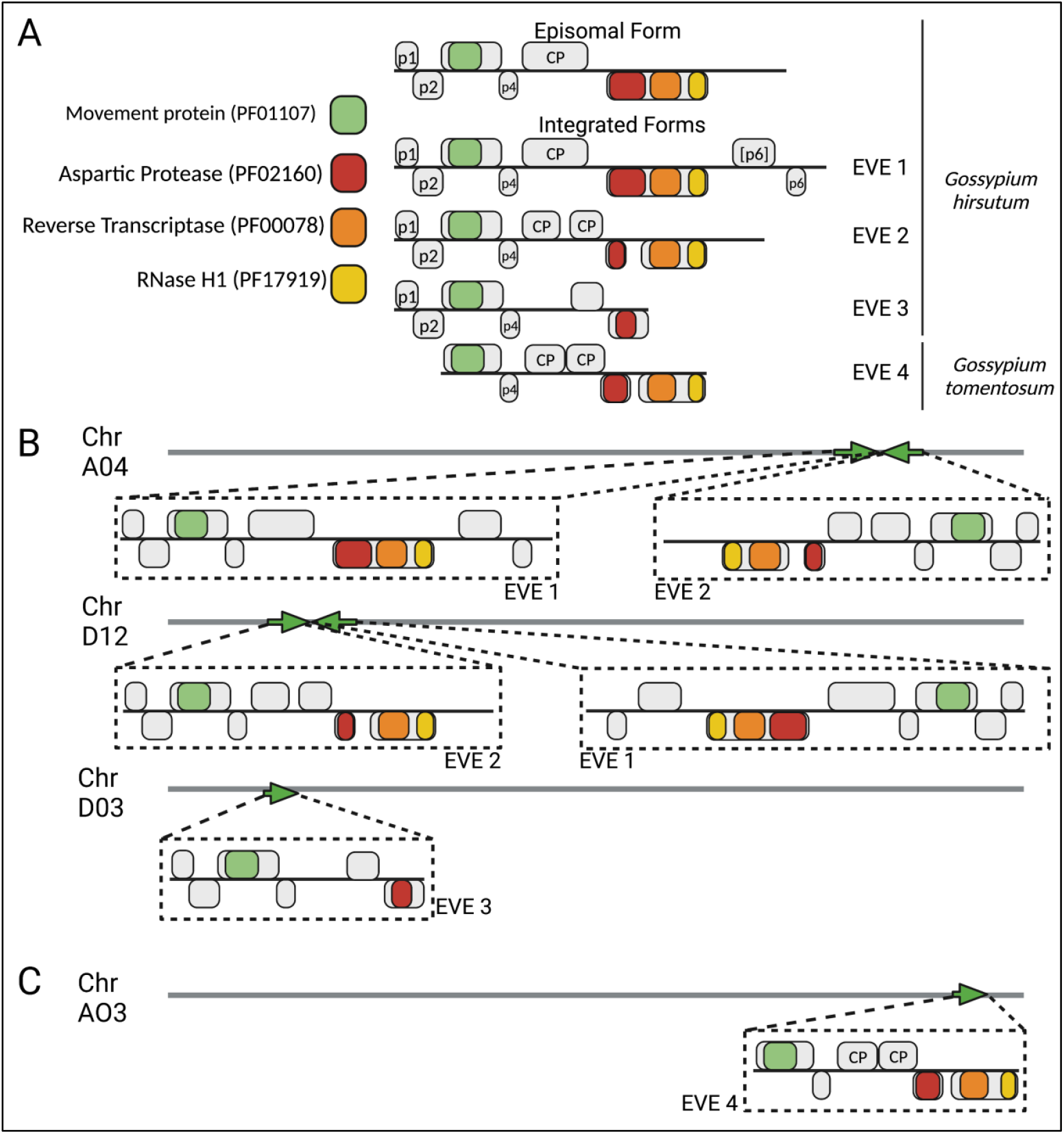
Illustration representing endogenous virus elements (EVEs) in *Gossypium* spp. genomes that likely originated from infection by ancestor of cotton virus A (CotV-A). EVEs were detected *in silico* in *G. hirsutum* and *G. tomentosum* chromosome-level genomes available in the CottonGen database. A) EVEs 1-4 present unique, structurally different gene organizations compared to CotV-A. B) EVE 1 and 2 in *G. hirsutum* were found as inverted tandem repeats in chromosomes A04 (11 out of 13 genomes) and D12 (1 out of 13 genomes). EVE 3 was found in chromosome D03 (13 out of 13 genomes) as a single integration event and was considerably shorter than the episomal CotV-A genome. C) EVE 4 found in *G. tomentosum* that was found as a single integration event in chromosome A03.

To infer the evolutionary relationships between CotV-A and the EVEs, phylogenetic analyses were performed as detailed above using the MP and peptidase sequences, which are the only proteins with caulimovirid homologs present among the episomal virus and the EVEs. For both proteins, phylogenetic analyses showed that EVE4 is more closely related to an ancestor virus of CotV-A, than the other EVEs and CotV-A (Figure S1). Together with the absence of an EVE 4 homolog in chromosome A03 of *G. hirsutum*, these results support the hypothesis that an ancestor of CotV-A infected both *Gossypium* species after speciation took place, and at different time frames, with the infection and subsequent viral element integration into *G. hirsutum* likely being more recent. Intriguingly, the phylogenic analysis of the MP shows that CotV-A and its EVEs are more closely related to soymoviruses than to caulimoviruses (Figure S1A), suggesting that CotV-A could be a recombinant virus resulting from co-infection of a soymovirus and a caulimovirus.

Due to the unique attributes that CotV-A presents, such as the lack of ORFs presenting homology to proteins involved in aphid transmission (caulimovirus P2 and P3) and viroplasm formation (caulimovirus P6), further molecular and biological studies are required to understand if it is pathogenic in cotton. In addition, the EVEs in the genomes of *G. hirsutum* and *G. tomentosum* should be further corroborated because the analysis was performed using publicly available genomic data, which may represent incomplete or inaccurate cotton genome assemblies. It will also be interesting to determine whether CotV-A is aphid transmissible, and forms viroplasms akin to other caulimoviruses. Moreover, further experiments are needed to determine if abiotic factors such as heat and drought, and/or biotic factors such as insect feeding and/or CLRDV or other virus co-infection, trigger the formation of the episomal form of CotV-A from the EVEs.

## Acknowledgements and Conflicts of Interest

This material is based upon work supported by the National Science Foundation Graduate Research Fellowship Program under No. Grant DGE–2139899. Any opinions, findings, and conclusions or recommendations expressed in this material are those of the authors and do not necessarily reflect the views of the National Science Foundation. The research was funded by NIFA CARE grant 8062-22410-007-014-I, USDA ARS project number 8062-22410-007-000-D, and Cotton Incorporated grant 20-214/58-8062-0-003. We are grateful to members of the Heck Lab for helpful discussion, and Chad Thomas, Lisa Scanlon and Julie Bojanowski for their help in maintaining the cotton plants used in the study. We are grateful to Jodi Scheffler from the Crop Genetics Research Unit (USDA Agricultural Research Service) for her helpful and invaluable assistance in plant and sample collection.

