## Supplementary for "Characterization of cotton virus A, a novel and distinct member of the genus *Caulimovirus* with endogenous viral elements in *Gossypium* spp"

**Supplementary Table S1.** Primer sequences used in this study for detection of the movement protein (MP) gene, and genome amplification of cotton virus A (CotV-A).

| Purpose | Primer Name | Sequence (5' - 3') | Product Size (bp) |
|---|---|---|---|
| Diagnostics MP | Cot-Caulimo-MP-F | GGACGACTCGAAGGAAACTTAGG | 474 |
| | Cot-Caulimo-MP-R | ACTAGAAGGGTGCTCTACTGGTA | |
| Genome amplification | Cot-Caulimo-F1 | ACGAATGGAGACAGGATTGAGG | 1315 |
| | Cot-Caulimo-R1 | AGAATTCCACCATGCTCGTACA | |
| | Cot-Caulimo-F2 | GGTTGGTACCGTCAAATGAGC | 1545 |
| | Cot-Caulimo-R2 | CGGGTCGATGCCAACTGTA | |
| | Cot-Caulimo-F3 | GCAAGGAACGTTGTTTAAGGTCC | 1566 |
| | Cot-Caulimo-R3 | GCTTTCTTTACAGCGAGCCA | |
| | Cot-Caulimo-F4 | AGCCAAGAATATTGGTCTGGTGT | 1189 |
| | Cot-Caulimo-R4 | CTCGATCCAGACCGAAATGTCT | |
| | Cot-Caulimo-F5 | GTGGGTAGTAGAGTGGGTCG | 2239 |
| | Cot-Caulimo-R5 | TGCTGAATGGGTAATCATCCTGA | |

**Supplementary Table S2.** Nucleotide and amino acid sequence identity (%) of cotton virus A (CotV-A) with other members of the family *Caulimoviridae.* Accession numbers for established *Caulimoviridae* members were obtained from the International Committee on Taxonomy of Viruses (ICTV) database. Plant-associated caulimovirus 1 and grapevine pararetrovirus are not officially classified, but are closely related to CotV-A.

| **Virus Name** | **Accession number** | **Max Score** | **Query coverage (%)** | **Whole genome nt Identity (%)** | **Predicted Protein Sequence Identity (%)** | | | | | |
|---|---|---|---|---|---|---|---|---|---|---|
| | | | | | **ORF1** | **ORF2** | **ORF3** | **ORF4** | **ORF5** | **ORF6** |
| Grapevine pararetrovirus (GPRV) | OP886324 | 365 | 22 | 66.74 | - | - | - | - | 33.57 | 54.40 |
| Figwort mosaic virus (FMV) | X06166 | 209 | 16 | 64.85 | - | - | 31.98 | - | 25.22 | 52.38 |
| Soybean Putnam virus (SPuV) | JQ926983 | 205 | 16 | 64.83 | - | - | 35.39 | - | 25.65 | 49.27 |
| Plant-associated caulimovirus 1 (PaCaV1) | OL472131 | 172 | 13 | 69.05 | - | - | 32.44 | - | 37.43 | 52.04 |
| Lamium leaf distortion virus (LLDV) | EU554423 | 153 | 6 | 68.32 | - | - | 31.98 | - | 27.50 | 49.35 |
| Angelica bushy stunt virus (AnBSV) | KU508800 | 148 | 10 | 65.60 | - | - | 31.55 | - | - | 47.54 |
| Dahlia mosaic virus (DMV) | JX272320 | 121 | 74 | 66.67 | - | - | 32.07 | - | 26.46 | 51.64 |
| Mirabilis mosaic virus (MMV) | AF454635 | 120 | 19 | 62.92 | - | - | 29.96 | - | - | 49.92 |
| Carnation etched ring virus (CERV) | X04658 | 107 | 10 | 63.75 | - | - | 31.86 | - | 28.00 | 48.77 |
| Strawberry vein banding virus (SBVB) | X97304 | 103 | 6 | 66.96 | - | - | 33.63 | - | 24.63 | 49.02 |
| Cauliflower mosaic virus (CaMV) | V00141 | 91.5 | 12 | 63.30 | - | - | 32.53 | - | 27.11 | 50.32 |
| Atractylodes mild mottle virus (AMMV) | KR080327 | 86.9 | 11 | 66.41 | - | - | 33.69 | - | 23.15 | 47.13 |
| Horseradish latent virus (HLV) | JX429923 | 54.5 | 2 | 80.70 | - | - | 31.28 | - | 24.05 | 47.71 |

**Supplementary Table S3.** Nucleotide and amino acid sequence identities (%) of cotton virus A (CotV-A) compared with the endogenous virus elements (EVEs) found in *Gossypium hirsutum* on chromosomes A04 (EVEs 1 and 2), D12 (EVEs 1 and 2), and D03 (EVE 3), and in *G. tomentosum* on chromosome A03 (EVE 4).

| ***Gossypium* species** | **Endogenous virus element (EVE)** | **Length (bp)** | **Query coverage (%)** | **Whole genome nt identity (%)** | **Predicted Protein Sequence Identity (%)** | | | | | |
|---|---|---|---|---|---|---|---|---|---|---|
| | | | | | **ORF1** | **ORF2** | **ORF3** | **ORF4** | **ORF5** | **ORF6** |
| *G. hirsutum* | EVE 1 | 8004 | 83.2 | 99.2 | 99.2 | 98 | 98 | 98.2 | 98.3 | 97.1 |
| *G. hirsutum* | EVE 2 | 7036 | 94 | 99.6 | 100 | 100 | 100 | 100 | 89.3 | 84.5 |
| *G. hirsutum* | EVE 3 | 4760 | 60.5 | 79.6 | 83.6 | 85.9 | 94.1 | 71.6 | - | 54.3[a] |
| *G. tomentosum* | EVE 4 | 4924 | 66.4 | 79.4 | - | - | 88.8 | 75.2 | 71.1 | 85 |

[a] Protein identity corresponds to the peptidase homolog region.

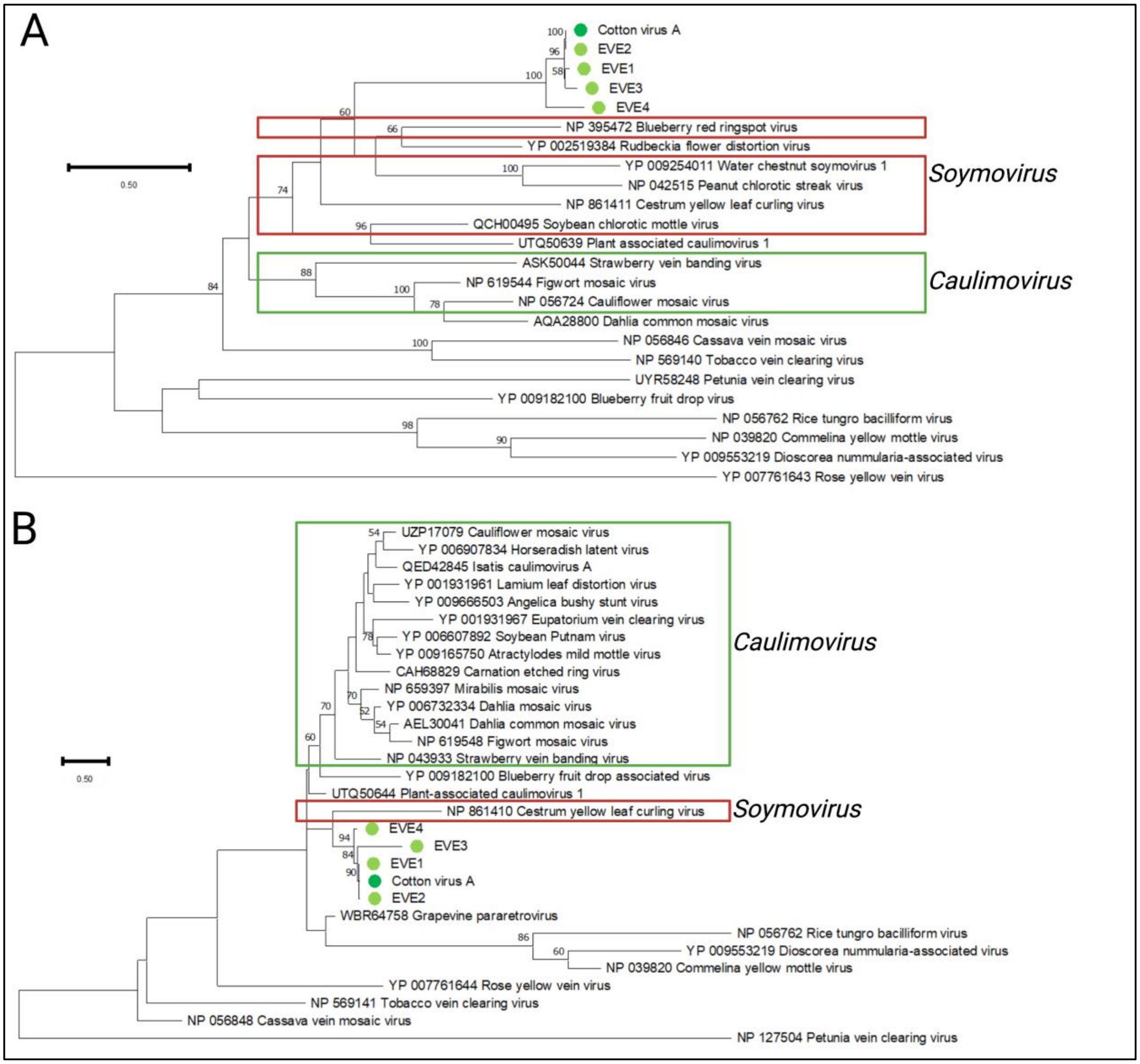


**Figure S1.** Phylogenetic relationships of movement protein (MP) (A) and peptidase (B) protein sequences of cotton virus A (CotV-A) and its endogenous virus elements (EVEs) to other members of the family *Caulimoviridae*. The maximum-likelihood method with the best model of protein evolution (LG + G) was used with 1,000 bootstrap pseudoreplicates as percentage values for branch support. Predicted amino acid sequences were used, and the respective GenBank accession number is shown with each virus name. The alignment was generated using MUSCLE and implemented in MEGA v.11.0.13. The dark green circle indicates CotV-A, while the light green circles indicate EVEs.

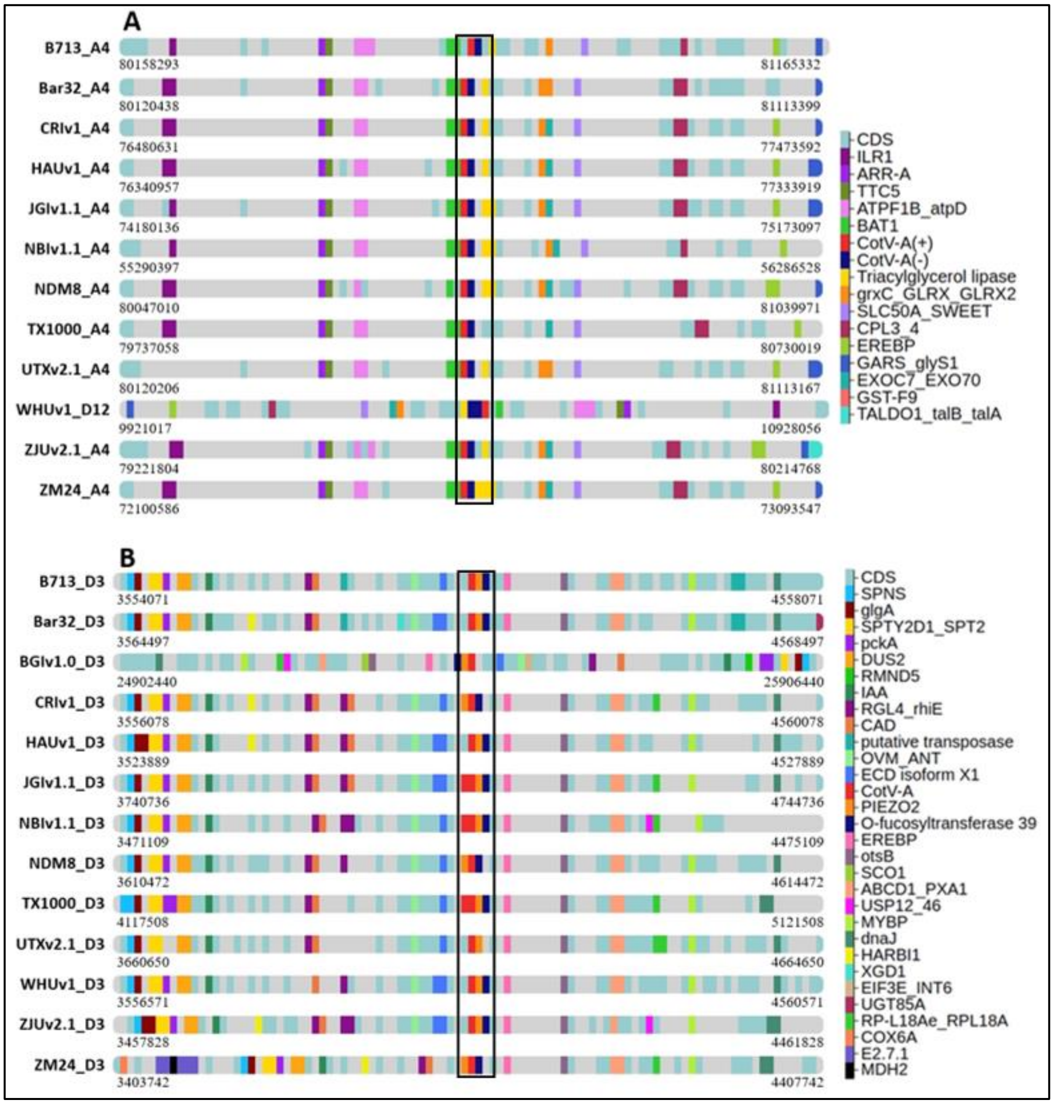


**Figure S2.** Windows showing 1 Mbp of genomic sequence adjacent to the endogenous virus element (EVE) integration sites. Coding sequences (CDS) on *Gossypium hirsutum* chromosomes A4, D12, and D3. CDS are unannotated genes, and numbers below each chromosome window are the coordinates within each designated chromosome. A) EVEs 1 and 2 in chromosomes A4 and D12. B) EVE 3 in chromosome D03. The assigned names for the 13 *G. hirsutum* genomes used in this study that were retrieved from cottongen database are as follows: 'B713' genome NSF_v1 (B713), 'Bar32' genome NSF_v1 (Bar32), 'TM-1' genome CGP-BGI_v1 (BGIv1.0), 'TM-1' genome CRI_v1 (CRIv1), 'TM-1' genome HAU_v1 (HAUv1), 'TM-1' genome UTX-JGI-Interim-release_v1.1 (JGIv1.1), 'TM-1' genome NAU-NBI_v1.1 (NBIv1.1), 'NDM8' genome HEAU_v1 (NDM8), 'TX-1000' genome CRI_v1 (TX1000), 'TM-1' genome UTX_v2.1 (UTXv2.1), 'TM-1' genome WHU_v1 (WHUv1), 'TM-1' genome ZJU-improved_v2.1_a1 (ZJUv2.1), 'ZM24' genome CRI_v1 (ZM24).

**Figure S2A gene names:** IAA-amino acid hydrolase (ILR1), two-component response regulator ARR-A family (ARR-A), tetratricopeptide repeat protein 5 (TTC5), F-type H+/Na+-transporting ATPase subunit beta (ATPF1B_atpD), Spliceosome RNA helicase

BAT1 (BAT1), glutaredoxin 3 (grxC_GLRX_GLRX2), solute carrier family 50 sugar transporter (SLC50A_SWEET), RNA polymerase II C-terminal domain phosphatase-like 3/4 (CPL3_4), EREBP-like factor (EREBP), glycyl-tRNA synthetase (GARS_glyS1), exocyst complex component 7 (EXOC7_EXO70), GSTF9 glutathione S-transferase PHI 9 (GSTF9), transaldolase (TALDO1_talB_talA).

**Figure S2B gene names:** MFS transporter_Spinster family_sphingosine-1-phosphate transporter (SPNS), starch synthase (glgA), protein SPT2 (SPTY2D1_SPT2), phosphoenolpyruvate carboxykinase (ATP (pckA), tRNA-dihydrouridine synthase 2 (DUS2), E3 ubiquitin-protein transferase RMND5 (RMND5), auxin-responsive protein IAA (IAA), rhamnogalacturonan endolyase (RGL4_rhiE), cinnamyl-alcohol dehydrogenase (CAD), Protein ecdysoneless isoform X1 (ECD isoform X1), Piezo type mechanosensitive ion channel component 2 (PIEZO2), rhamnogalacturonan I rhamnosyltransferase (RRT), EREBP-like factor (EREBP), trehalose-6-phosphate phosphatase (otsB), protein SCO1 (SCO1), ATP-binding cassette_subfamily D (ALD (ABCD1_PXA1), ubiquitin carboxyl-terminal hydrolase 12/46 (USP12_46), transcription factor MYB_plant (MYBP), molecular chaperone DnaJ (dnaJ), nuclease HARBI1 (HARBI1), xylogalacturonan beta-1), 3-xylosyltransferase (XGD1), translation initiation factor 3 subunit E (EIF3E_INT6), UDP-glucosyltransferase 85A (UGT85A), large subunit ribosomal protein L18Ae (RP-L18Ae_RPL18A), cytochrome c oxidase subunit 6a (COX6A), kinase (E2.7.1), malate dehydrogenase (MDH2).
